# Caffeinated soda intake in children is associated with neurobehavioral risk factors for substance misuse

**DOI:** 10.1101/2021.02.17.431718

**Authors:** Mina Kwon, Hyeonjin Kim, Jaeyeong Yang, Yoseph Lee, Jihyun K. Hur, Tae-Ho Lee, James M. Bjork, Woo-Young Ahn

## Abstract

**Background and Objectives:** Use of psychotropic substances in childhood has been associated with both impulsivity and other manifestations of poor executive function as well as escalation over time to use of progressively stronger substances. However, how this relationship may start in earlier childhood has not been well explored. Here, we investigated the neurobehavioral correlates of daily caffeinated soda consumption in preadolescent children and examined whether caffeinated soda intake is associated with a higher risk of subsequent alcohol initiation.

**Methods:** Using Adolescent Brain Cognitive Development study data, we first investigated cross-sectional relationships between frequent caffeinated soda intake and well-known risk factors of substance misuse: impaired working memory, high impulsivity, and aberrant reward processing. We then examined whether caffeinated soda intake at baseline predicts more alcohol sipping at 12 months follow-up using a machine learning algorithm.

**Results:** Daily consumption of caffeinated soda was cross-sectionally associated with neurobehavioral risk factors for substance misuse such as higher impulsivity scores and lower working memory performance. Furthermore, caffeinated soda intake predicted greater alcohol sipping after 12 months even after controlling for rates of baseline alcohol sipping; children who drink caffeinated soda daily are twice as likely to start sipping alcohol after one year compared to those who do not drink caffeinated soda at all.

**Conclusions:** These findings suggest that previous linkages between caffeine and substance use in adolescence also extend to younger initiation, and may stem from core neurocognitive features thought conducive to substance initiation.

---

Soft drinks are commonly consumed even by children, and a vast majority of sodas contain caffeine (Temple, 2009, 2018). Moreover, caffeinated sodas typically contain sweeteners such as high-fructose corn syrup, which can affect neurocognitive function and cause physical side effects, for an instance, by affecting microbiome (Ettinger, 2022). Not surprisingly, the consequences of excessive consumption of both sugar and caffeine have been well documented (Ooi et al., 2022; Porciúncula et al., 2013; Temple et al., 2017), including a strong association between caffeinated beverage consumption in adolescence and future substance use (Arria et al., 2011; Barrense-Dias et al., 2016; Kristjansson et al., 2018; Leal & Jackson, 2019; Marmorstein, 2018; Miyake & Marmorstein, 2015). In prospective studies tracking the effects of substance use, the percentage of regular energy drink users who became alcohol or marijuana users after 1– 2 years was approximately five times higher than that of non-energy drink users (Leal & Jackson, 2019; Marmorstein, 2018; Miyake & Marmorstein, 2015). Others have shown that coffee or energy drink consumption in adolescents or young adults significantly predicts future substance use, such as tobacco and alcohol use (Arria et al., 2011; Barrense-Dias et al., 2016; Kristjansson et al., 2018, 2022; Marmorstein, 2018).

In light of the potential for a problematic progression from drinking caffeinated beverages to using harder substances, there is a critical need to investigate aspects of this progression as early as possible. As adolescence is the most common period for initiating substance use and an earlier onset of substance use predicts greater addiction severity (Chambers et al., 2003; Jordan & Andersen, 2017), examining these relationships in preadolescence is crucial in that caffeinated soda intake in childhood could provide useful predictive information on future substance use. In preadolescent children, caffeinated soda is the most preferred and accessible mode of caffeine intake (Temple, 2009, 2018).

Only a handful of studies, however, have directly examined the effects of frequent caffeinated beverage consumption in preadolescent children, due to their lower rates of consumption (0.4% for coffee and < 0.1% for energy drinks in 9–10-year-old children) (Lisdahl et al., 2021). Most of the previous studies examining the association between caffeine consumption and later substance use have focused primarily on adolescents who drink multiple caffeinated beverages on a daily basis (Temple, 2009, 2018). Moreover, while a few existing studies have examined behavioral risk factors associated with caffeinated soda intake, they have not assessed neural underpinnings of such risk factors (James et al., 2011; Miyake & Marmorstein, 2015; Solnick & Hemenway, 2013; Suglia et al., 2013).

Here, we addressed the unanswered question of whether frequent consumption of caffeinated soda in preadolescent children indicates a higher risk of future alcohol experimentation, using data from the Adolescent Brain Cognitive Development (ABCD) Study (Bjork et al., 2017). In addition, we wished to explore the potential neurobehavioral mechanisms of such relationships. In light of previous linkages between activity in neurocircuits germane to motivation and inhibition (as detected from functional magnetic resonance imaging (fMRI)) and substance use in adolescents (Lees et al., 2021), we examined the relationship between daily caffeinated soda intake and both behavioral and neuroimaging markers of neurobehavioral risk factors for substance use disorders (SUDs) in children. These include impaired working memory (WM), high impulsivity, and altered mesolimbic reward processing (**Fig. 1A**). These are the three primary neurocircuit functions targeted in the ABCD study due to their significant implications for addiction (see Casey et al. (2018) for more detail). We applied a machine learning approach (a least absolute shrinkage and selection operator (LASSO) regression; Tibshirani, 1996) to the measures to identify multivariate risk factors for SUDs associated with daily caffeinated soda intake and select features highly associated with caffeinated soda intake (Volkow et al., 2015). Using the same approach, we then examined whether caffeinated soda intake could predict future alcohol sipping, as (1) alcohol sipping has been reported to predict future alcohol abuse (Jackson et al., 2015; Watts et al., 2021), and (2) alcohol sipping is the most common gateway behavior toward other substances of abuse (Barry et al., 2016). Based on the previous literature suggesting that caffeinated beverage consumption predicts future substance use (Arria et al., 2011; Barrense-Dias et al., 2016; Leal & Jackson, 2019; Marmorstein, 2018; Miyake & Marmorstein, 2015), we hypothesized that daily caffeinated soda intake in the baseline ABCD assessment (children age 9-10) would predict future alcohol sipping at the year 1 follow up (**Fig. 1B**). Taken together, we aimed to elucidate the potential neurobehavioral underpinnings of frequent caffeinated soda intake during childhood.

**Figure 1.**
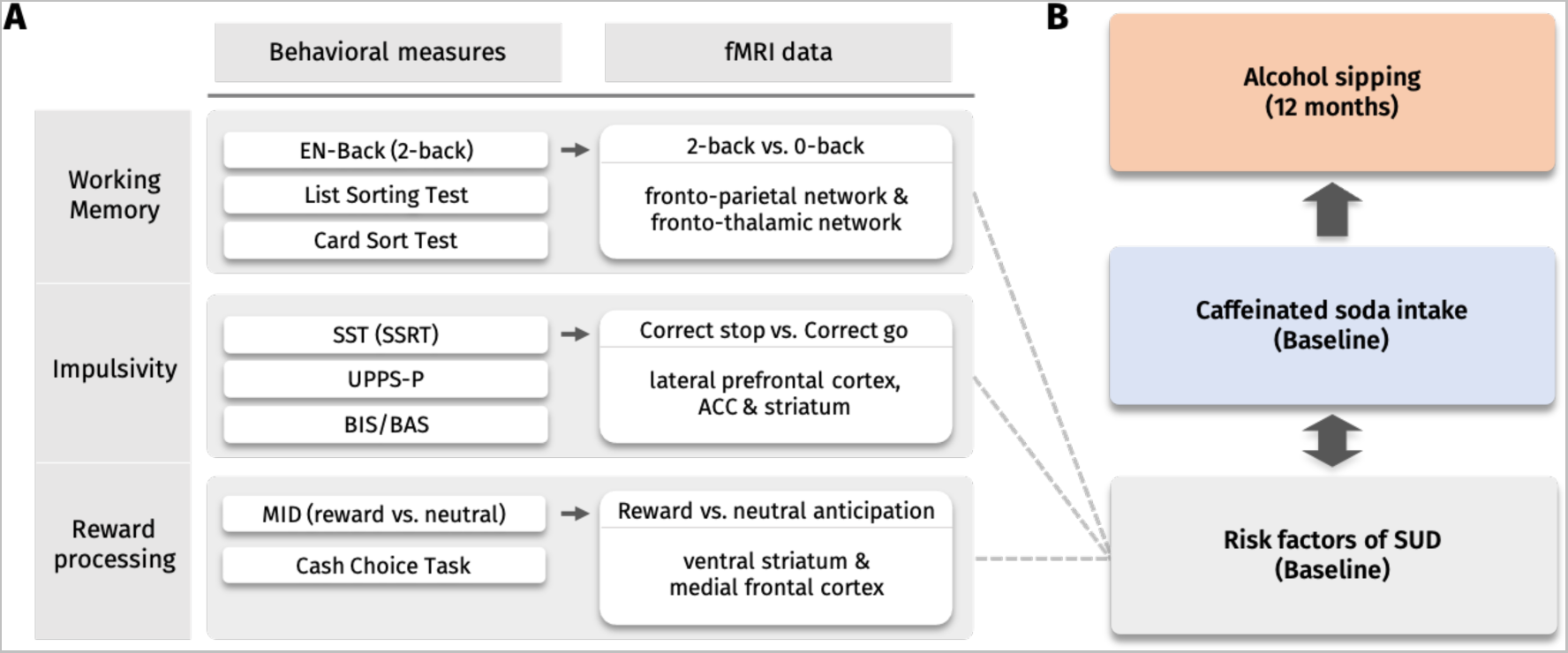
An overview of the analytical approach. **A.** The assessments used to capture the three main categories of neurobehavioral risk factors: working memory, impulsivity, and reward processing. Working memory was measured from the performance of list sorting test, card sort test, and 2-back in the emotional N-Back Task (EN-back) performed inside a functional magnetic resonance imaging (fMRI) scanner. The fMRI data from the EN-back task were analyzed based on the contrast of 2-back versus 0-back, using regions in the frontoparietal and fronto-thalamic network as the regions of interest (ROIs). Impulsivity was measured by Urgency-Premeditation-Perseverance-Sensation Seeking-Positive Urgency (UPPS-P), and Behavioral Inhibition System/ Behavioral Activation System (BIS/BAS), and the stop signal reaction time (SSRT) during the stop signal task (SST) performed inside the fMRI scanner. The fMRI data from the SST were analyzed based on the contrast of correct stop versus correct go in lateral prefrontal cortex, anterior cingulate cortex (ACC), and striatum ROIs. Reward processing was measured behaviorally by the cash choice task, and also by comparing the success rate of reward versus neutral conditions during the monetary incentive delay (MID) task performed inside the fMRI scanner. The fMRI data from the MID task were analyzed based on the contrast of reward anticipation versus neutral anticipation, using regions in the ventral striatum and medial frontal cortex as the ROIs. The ROIs were selected based on the Destrieux atlas (Destrieux et al., 2010). See “**Methods**” for more details on the variables. **B.** Diagram of the research aims. First, the associations between the risk factors for substance use disorders (SUDs) and caffeinated soda intake were examined cross-sectionally using the baseline ABCD data. Then, we assessed whether caffeinated soda intake would predict alcohol sipping after 12 months while controlling other risk factors of SUDs.

## Methods

### Participants

The ABCD study collected data from 11,878 children aged 9–10 years. The participants were recruited via school systems from 21 different sites in the United States. The Institutional Review Board (IRB) at the University of California, San Diego, approved all the research protocol of the ABCD study at most sites per reliance agreement, with local IRB approval at the remaining sites (Auchter et al., 2018). All participants provided written assent, and their parents or guardians provided written consent (Auchter et al., 2018). More information about the recruitment and study design is available in Garavan et al. (2018). Further details of the demographic, physical, and mental health assessments are described in Barch et al., (2018).

Out of 11,878 children from ABCD release 2.0, we excluded those with any missing data regarding either caffeinated soda intake, neurobehavioral risk factors for SUDs, future alcohol sipping, and confounding variables (see **Measures**). We also excluded outliers based on the measures of caffeinated soda intake with the cutoff larger/smaller than the mean ±5 standard deviations. Of these, 147 reported drinking more than 7 cans of caffeinated soda per week (daily-drinkers) and 1,945 reported not drinking a single can (non-drinkers) in the past 6 months. Therefore, we included 2,092 participants for the main analyses (see **Fig. S1** for a flowchart of the selection process). To examine the potential for selection bias, we compared the baseline characteristics among children who were included and excluded in the analyses (**Table S1**).

### Measures

#### Caffeinated soda intake

Caffeinated soda intake was assessed from self-report of the participants in response to the question, “How many drinks of the following beverages have you had per week in the past 6 months? – soda with caffeine (Mountain Dew, Jolt, Coke, Pepsi, Dr. Pepper, Barq’s root beer)”. The participants who reported consuming more than 7 cans per week (i.e., corresponds to more than 1 can per day in average) were allocated to daily-drinking group (*N*=147), and those who reported consuming 0 can per week were allocated to the non-drinking group (*N*=1,945). See **Table S2** for a comparison of the (average) daily-drinkers and non-drinkers on the following behavioral measures, and see **Table S3** for a comparison of the neural measures.

#### Neurobehavioral risk factors for SUDs

As shown in **Fig. 1A**, we focused on three well-known neurobehavioral risk factors for SUDs: poor WM, increased impulsivity, and aberrant reward processing. These constructs were collectively measured by self-report surveys, behavioral tasks, and fMRI, respectively, as described below. Spatial regions of interest (ROIs) for each task were restricted *a priori* to the canonical activations initially reported using the ABCD samples (Casey et al., 2018). The curated data used were based on the Destrieux atlas (Destrieux et al., 2010).

To obtain behavioral measures of WM, we used the List Sorting Working Memory Test (list sorting test; Tulsky et al., 2013) and Dimensional Change Card sort Test (card sort test; Tulsky et al., 2013), as well as response accuracy during the 2-back condition of the emotional N-back (EN-back) task (Cohen et al., 2015) (see **Supplementary materials**). The EN-back task was conducted inside the MRI scanner. The contrast of “2-back versus 0-back” was used for the fMRI analysis, and the ROIs were selected based on activation maxima of the contrast in the initial subset of participants (Casey et al., 2018): the rostral middle frontal gyrus (MFG), caudal MFG, inferior frontal gyrus (IFG; pars triangularis and pars orbitalis), lateral orbitofrontal cortex (OFC), superior parietal lobule and inferior parietal lobule (IPL) in the frontoparietal network, and the caudate nucleus, putamen, nucleus accumbens (NAc), rostral anterior cingulate cortex (ACC), caudal ACC, thalamus proper, ventral diencephalon, amygdala, and hippocampus in the fronto-thalamic network.

Due to the multi-faceted construct of impulsivity, wherein laboratory and self-report assessments of impulsivity are thought to capture different components (Sharma et al., 2014), we analyzed each of self-report trait-like impulsivity as well as rapid-response and decision-based impulsivity. Trait impulsivity was measured using the short form Urgency–Premeditation–Perseverance–Sensation Seeking–Positive Urgency (UPPS-P) impulsive behavior scale for children (the 20-item short version for youths; Barch et al., 2018) and a parental report of the ABCD Youth Behavioral Inhibition System/ Behavioral Activation System (BIS/BAS; Carver & White, 1994). The BAS is related to goal-directed efforts, such as motor activation in response to an impending reward, while the BIS is engaged when inhibition toward a goal occurs, such as the avoidance of punishment (Carver & White, 1994). Excessive behavioral inhibition is associated with depression and anxiety, while excessive behavioral activation is associated with impulsive behaviors, compulsive behaviors, substance misuse, and aggression (Newman et al., 2005). To identify the neurocircuit correlates of impulsivity, participants completed the stop signal task (SST; Logan et al., 1984) during fMRI. The SST measures impulsivity related to impulse control or response inhibition when performing an action. The contrast of “correct stop versus correct go” was used for the fMRI analysis. ROIs were the lateral prefrontal cortex (rostral MFG, pars orbitalis in IFG, and lateral OFC), rostral and caudal ACC, and striatum (caudate, putamen, and NAc). These areas have been associated with impulsivity and impulse control (Aron et al., 2014; Casey et al., 2018). See **Supplementary materials** for more details of the SST.

Reward processing is closely linked to impulsivity, as impulsive people often show immediate reward-seeking behavior (Zuckerman, 2001). For our offline metric of reward sensitivity, we used the Cash Choice Task (Luciana et al., 2018), which assesses willingness to delay gratification (see **Supplementary materials**). Previous studies have suggested that people with SUDs exhibit hyper-responsiveness of mesolimbic motivational neurocircuitry toward drug-related cues (Bechara et al., 2019) while showing blunted responsiveness toward cues for a non-drug reward (e.g., monetary reward) (Luijten et al., 2017). The monetary incentive delay (MID) task (Knutson et al., 2000) is widely used to measure the neural correlates of the anticipation of monetary rewards and losses. The contrast of “reward versus neutral” at the cue onset, which reflects reward anticipation, was used for the fMRI analysis, with the ROIs in the ventral striatum (i.e., NAc), rostral ACC, and medial OFC in the medial prefrontal cortex, which play a key role in reward processing, particularly in reward anticipation (Bartra et al., 2013). See **Supplementary materials** for more details about the MID task.

#### Future alcohol sipping

Alcohol sipping was measured by self-report from the participants using the iSay Sip Inventory, which was performed once a year and asked only in children who had heard of alcohol (see Lisdahl et al. (2018) for more details). From the year 1 data (data release 3.0), we used a binary response to a single question asking if the participants had sipped alcohol outside of a religious ceremony. We considered those who had not heard of alcohol as having yet to sip alcohol (i.e., no alcohol sipping).

#### Confounding variables

In light of the potential for socioeconomic and structural factors to influence health behaviors, we included the following variables as confounding variables in the data analysis: socioeconomic status (SES), family history of substance use, parental monitoring, sleep deprivation, externalizing/internalizing symptomatology, data collection sites, and type of MRI scanner. See **Supplementary materials** for more detail.

### Analysis

#### Caffeinated soda intake and risk factors for SUDs

To identify a multivariate profile of risk factors for caffeinated soda consumption, we performed a binomial LASSO regression analysis (Tibshirani, 1996), a machine learning algorithm, to distinguish the daily soda-drinking group (*N*=147) from the non-drinking group (*N*=1,945). We used all of the risk factors and confounding variables as input (i.e., predictors) in the LASSO-based prediction model to classify each individual into the daily-drinking group (coded as “1”) or non-drinking group (coded as “0). More specifically, candidate predictors included every measure for the three cognitive factors as well as other control variables. As LASSO regression offers feature selection (Tibshirani, 1996), we aimed to identify features that could differentiate the daily-drinking group from the non-drinking group. See **Supplementary materials** for more details on LASSO regression.

#### Caffeinated soda intake and future alcohol sipping

We performed a binomial LASSO regression analysis to test the association between daily caffeinated soda intake and alcohol sipping. We used all possible variables, including the three primary neurobehavioral risk factors (WM, impulsivity, and reward processing) along with the confounding variables as candidate predictors collected at baseline. Using these inputs (i.e., predictors), the LASSO-based model was used to classify individuals who reported alcohol sipping after 12 months (coded as “1”; *N*=90) and those who reported no alcohol sipping (coded as “0”; *N*=2,002). We aimed to select predictors that could distinguish individuals who reported alcohol sipping from those who did not. All of the procedures, except for the predictors and dependent variables, are identical to those in the former analysis (see “**Methods - *Caffeinated soda intake and risk factors for SUDs”*)**. Furthermore, we performed a chi-square test to compare the ratio of alcohol sipping after 12 months between the two groups. Lastly, we estimated the risk ratio of the daily-drinking group by dividing the cumulative incidence of alcohol sipping in the daily-drinking group by the cumulative incidence in the non-drinking group.

To rigorously examine the association between caffeinated soda intake and future alcohol sipping, we conducted three additional analyses. First, we used continuous measure for caffeinated soda intake instead of using the categorical measure of daily- and non-drinkers. Using data from 4,517 participants (see **Fig. S1**), we carried out the same binomial LASSO regression analysis mentioned earlier (**Fig. S3**). Second, we used only the behavioral measures of risk factors in predicting future alcohol sipping, to include as many participants as possible. Using data from 8,939 participants (see **Fig. S1**), we again performed the binomial LASSO regression analysis (**Fig. S4**). Lastly, using the sample of 8,939 participants, we ran a hierarchical logistic regression analysis to confirm whether continuous measures of caffeinated soda intake at baseline can predict alcohol sipping after 12 months (**Table. S4**). For more details, please refer to the **Supplementary materials**.

## Results

### Caffeinated soda intake and neurobehavioral risk factors for SUDs

Our first question was whether daily caffeinated soda intake is cross-sectionally associated with well-known neurobehavioral risk factors for SUDs. **Fig. 2A** shows the multivariate profiles from binomial LASSO regression analysis distinguishing the daily-drinking group from the non-drinking group. Family history of drug use and low parental education were the two strongest predictors of daily consumption, along with sex (being male), lack of sleep, low family income, racial identities (as African American), high body mass index (BMI), high externalizing behaviors, and low parental monitoring. **Fig. 2B** shows a receiver-operation characteristic (ROC) curve and its mean area under the curve (AUC) for classification of the daily-drinking and non-drinking groups. The mean AUC values were 0.80 and 0.72 for the training and test sets, respectively.

**Figure 2.**
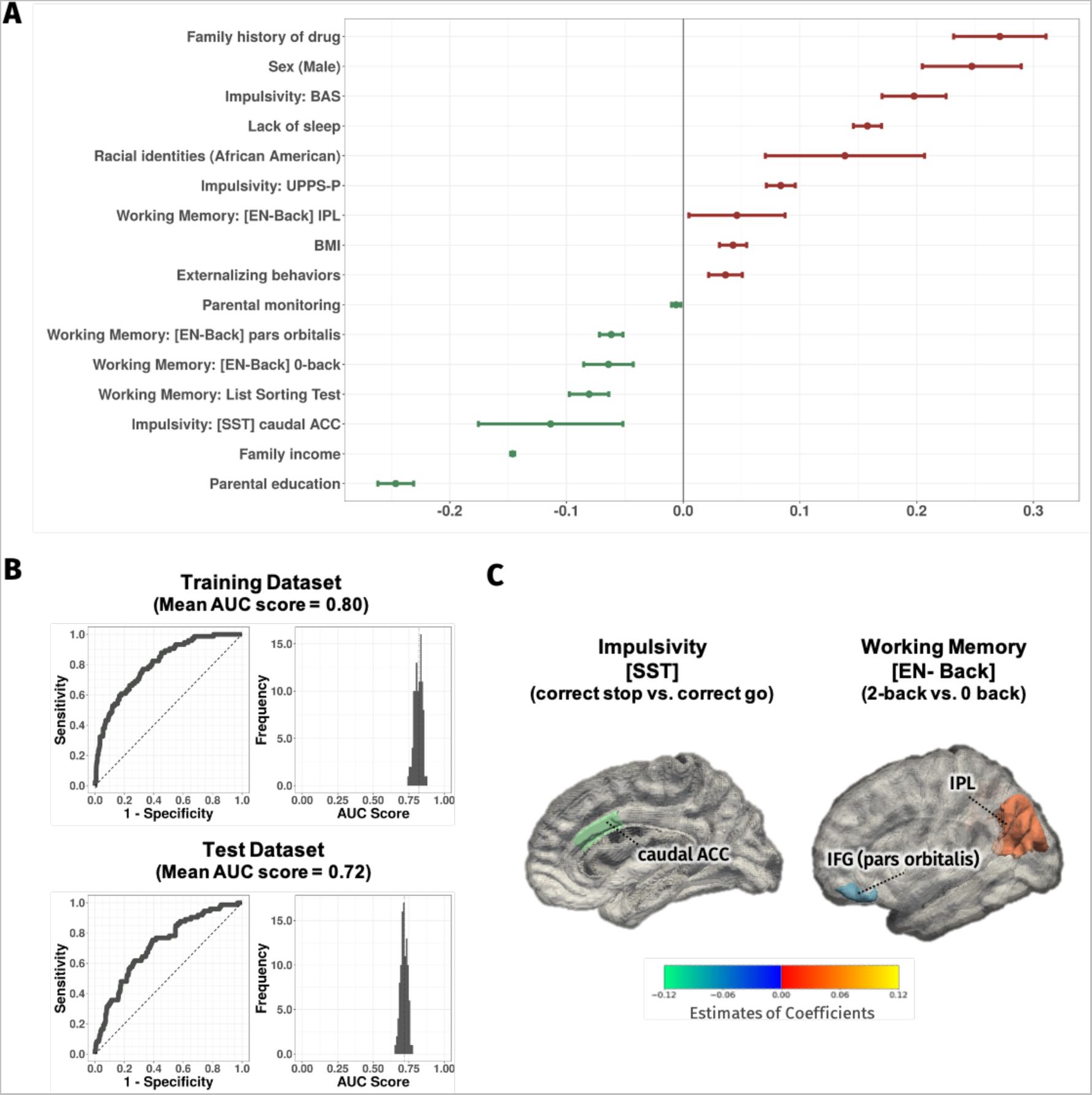
Results of binomial LASSO regression predicting daily caffeinated soda intake (daily-drinking group vs. non-drinking group) **A.** Estimates of coefficients of the survived variables (x axis: standardized coefficient estimates; y axis: predictor). The site variables were excluded for clarification (see **Supplementary Fig. S2** for results including all variables). Each dot indicates mean of the coefficient, positive coefficient in red and negative coefficient in green. Each error bar indicates 95% confidence interval. **B.** Distribution of the area under the curve (AUC) values (left) and a representative receiver-operation characteristic (ROC) curve (right) for the training and test datasets. **C.** Regions of interest (ROIs) identified as having significant estimates of coefficients during the stop signal task (SST) and emotional N-Back Task (EN-back). *Abbreviations*. LASSO, least absolute shrinkage and selection operator; BMI, body mass index; BAS, Behavioral Activation System; UPPS-P, Urgency-Premeditation-Perseverance-Sensation Seeking-Positive Urgency; IPL, Inferior Parietal Lobule; IFG, Inferior Frontal Gyrus; ACC, Anterior Cingulate Cortex.

Among our *a priori* candidate neurobehavioral risk factors for SUDs, high impulsivity measured from the BAS was most strongly associated with daily caffeinated soda intake (**Fig. 2C**). A higher UPPS-P score was also related to daily intake. Additionally, hypoactivation in the caudal ACC during the SST predicted the classification of the daily-drinking group. Among the WM measures, poor performance on the list sorting test and 0-back performance in the EN-back Task predicted daily intake of caffeinated soda. In addition, hypoactivation of the pars orbitalis of the IFG and greater activation of the IPL by working memory demands during the EN-back task (2-back vs. 0-back) also predicted daily caffeinated soda intake (**Fig. 2C)**. However, variables related to reward processing were not found to be predictors of daily-drinking group.

These results suggest that high impulsivity and low WM are significantly associated with daily caffeinated soda consumption. Along with some other demographic factors (family history of drug use, male sex, low SES, low parental monitoring, high externalizing behaviors, less sleep, and high BMI), the two neurobehavioral risk factors for SUDs distinguished the daily-drinking group from the non-drinking group.

### Caffeinated soda intake and future alcohol sipping

To address our second question of whether daily caffeinated soda intake predicts future alcohol sipping, we conducted binomial LASSO regression predicting alcohol sipping after 12 months, using all of the neurobehavioral risk factors and confounding variables collected at baseline. As shown in **Fig. 3A**, daily caffeinated soda intake was identified as one of the predictors that distinguished alcohol sipping after 12 months (mean estimate of coefficients=0.122, 95% confidence interval (CI)=[0.010, 0.235]), even after including alcohol sipping at baseline as a predictor. The mean AUC values of the predictive model were 0.90 and 0.72 for the training and test sets, respectively (**Fig. 3B**). Moreover, the ratio of alcohol sipping after 12 months was twice as high in the daily-drinking group compared to the non-drinking group (daily-drinking group: 0.082% (12 out of 147 children); non-drinking group: 0.040% (78 out of 1,945 children); X^2^ = 5.802, *p* = 0.016) (**Fig. 3C**). This finding suggests that the daily-drinking group had 2.04 times the risk of sipping alcohol compared to the non-drinking group. Our results were further supported by the binomial LASSO regression analyses with larger samples (**Fig. S4 & Fig. S5**) and hierarchical logistic regression analysis (**Table S4**) which can be found in the **Supplementary materials**.

**Figure 3.**
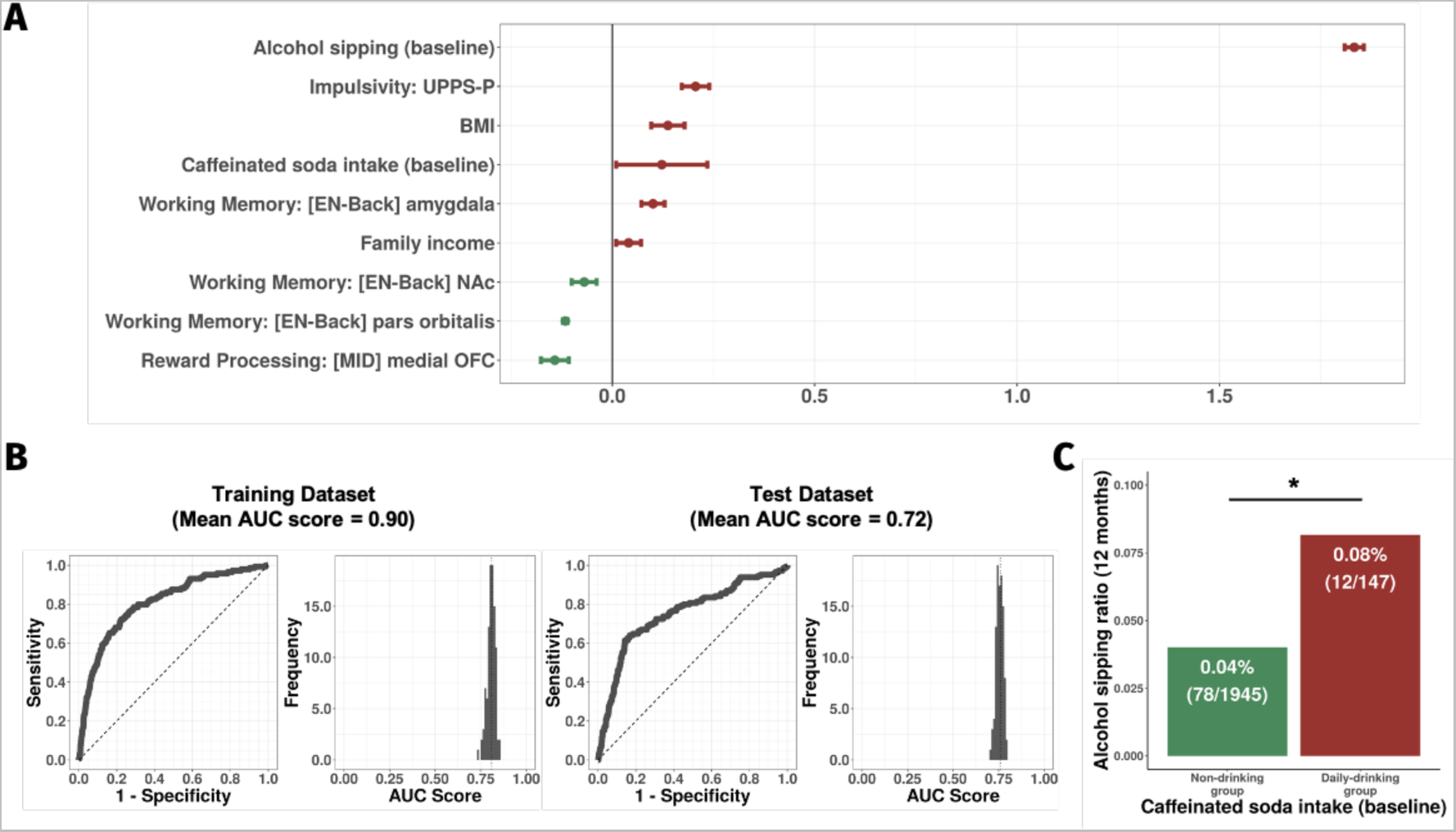
Results of binomial LASSO regression predicting alcohol sipping after 12 months. **A.** Estimates of coefficients of the survived variables (x axis: coefficient estimates; y axis: predictor). The site variables were excluded for clarification (see **Supplementary Fig. S3** for results including all variables). Each dot indicates mean of the coefficient, positive coefficient in red and negative coefficient in green. Each error bar indicates 95% confidence interval. **B.** Distribution of the area under the curve (AUC) values (left) and a representative receiver-operation characteristic (ROC) curve (right) for the training and test datasets. **C.** Rate of alcohol sipping at 12-month follow-up in each group. The asterisk indicates significance from the chi-square test (*p* < 0.05). *Abbreviations*. LASSO, least absolute shrinkage and selection operator; BMI, body mass index; UPPS-P, Urgency-Premeditation-Perseverance-Sensation Seeking-Positive Urgency; EN-Back, emotional N-Back Task; MID, Monetary Incentive Delay Task; NAc, Nucleus Accumbens; OFC, Orbitofrontal cortex.

## Discussion

In this study, we investigated whether caffeinated soda intake in preadolescent childhood is associated with a higher risk of alcohol experimentation in the future. Using the large dataset from the ABCD study, we first clarified cross-sectional relationships between daily caffeinated soda intake and well-known neurobehavioral risk factors for SUDs at study baseline; then evaluated whether caffeinated soda intake at baseline was predictive of alcohol sipping after 12 months. Our findings suggest that frequent consumption of caffeinated soda in children is closely related to previously-established neurobehavioral risk factors for SUDs and can predict future alcohol sipping.

Our machine learning approach partially supported our hypothesized association between caffeinated soda intake and the neurobehavioral risk factors for SUDs. Notably, high impulsivity scores and low WM performance each distinguished daily caffeinated soda drinkers from non-drinkers even after considering the effects of other confounding factors (**Fig. 2**). However, there was no significant association observed for mesolimbic recruitment by prospective rewards.

In relation to impulsivity, we found higher self-reported impulsivity in the daily-drinking group based on the UPPS-P and BAS scores along with altered activation in the ACC, a brain region implicated in cognitive control and impulsivity (Kerns et al., 2004; Shenhav et al., 2016). Reduced activities in the ACC during response inhibition or behavior-monitoring are commonly reported in children with attention deficit hyperactivity disorder (ADHD) (Cortese et al., 2012; Hart et al., 2013; McTeague et al., 2017) and individuals with SUDs (Luijten et al., 2014; Nestor et al., 2011; Yücel et al., 2007). Thus, hypoactivation of the ACC during response inhibition in daily-drinking group further seems to strengthen the association between daily soda consumption and elevated levels of impulsivity.

We also found WM impairments in the daily-drinking group on the list sorting test and the 0-back accuracy in the EN-back task, accompanied by hypoactivation in the IFG and hyperactivation in the IPL by working memory demands during the EN-back task. Prior studies have shown that greater activation in the prefrontal cortex is related to greater WM capacity (Casey et al., 2018; Owen et al., 2005), and increased activation in the IPL is associated with higher WM load (Baldo & Dronkers, 2006; Veltman et al., 2003). Taken together, these findings strongly suggest an association between daily-soda consumption and WM deficits.

In contrast, we did not find a significant association between neurocognitive measures of reward processing and daily caffeinated soda intake. Aberrant reward processing is a commonly observed neurocognitive feature in addiction (Zeng et al., 2023), with individuals either displaying increased or decreased sensitivity to reward (Berridge & Robinson, 2016; Blum et al., 2000; Demidenko et al., 2020; Robinson & Berridge, 1993). Altered reward processing has been also reported in children with a high risk of alcohol use, such as those with a family history of alcohol use problems (Bjork et al., 2008; Martz et al., 2022). One possibility for the lack of association is that the effect of reward processing variables on SUD may depend on population or context (Bjork, 2020), or may have been masked by the strong effect of family history of drug use, since our study controlled for the familial risk of alcohol and drug use. Additionally, it is worth considering that monetary rewards might have differential effects compared to drug rewards (Nestor & Ersche, 2023), which could explain the absence of significant associations. Consistent with our findings, a recent study comparing reward processing of alcohol dependent patients, first-degree relatives, and healthy controls could not find any significant group differences both in monetary reward and loss anticipation when controlling for age (Musial et al., 2023). Thus, further research is required to reconcile the mix findings related to reward processing in youth with a risk of addiction, perhaps by studying more narrowly defined subgroups and by examining both drug and non-drug rewards.

After examining the link between the neurobehavioral risk factors and caffeinated soda intake, we showed that frequent consumption of caffeinated soda predicted alcohol sipping after 12 months using LASSO regression (**Fig. 3**). Even after controlling for other well-established variables for alcohol sipping, including baseline alcohol sipping, caffeinated soda intake remained predictive of future alcohol sipping (**Fig. 3A**). Other survived predictors of future sipping include predictors for daily caffeinated soda intake itself (**Fig. 2A**), such as higher behavioral impulsivity score (i.e., UPPS-P), BMI, and hypoactivation of the IFG during the EN-back task. Conversely, reduced activity in the NAc and greater activity in the amygdala during the EN-back task, and reduced activity of the medial OFC during the MID task were associated with the participants who experienced alcohol sipping, but did not survive as predictors for daily caffeinated soda intake. Higher family income was associated with the participants who experienced alcohol sipping, while it was associated with the participants who do not drink caffeinated soda. Interestingly, some predictors of caffeinated soda intake were similar to the risk factors for SUDs (Jordan & Andersen, 2017), suggesting that caffeinated soda intake during childhood and SUDs share similar neurobehavioral vulnerabilities. Note that the majority of the cohort was substance-naïve (Lisdahl et al., 2018), and we evaluated alcohol sipping measures instead of substantial self-administration of alcohol. Thus, it would be informative to track the predictive ability of caffeinated soda and alcohol use and investigate changes in the direction of the predictors as the children get older.

Our findings suggest that caffeinated soda in children is predictive of substance use in the near future. The longitudinal associations between the use of more benign psychotropic substances early in life and the use of “harder” substances later in adolescence or emerging adulthood have been characterized for decades (Kandel, 1975), and have been attributed to two competing (but not mutually-exclusive) theories. The “gateway hypothesis” (Kandel & Yamaguchi, 2002) generally implies that exposure to the earlier-used substance itself, such as nicotine or cannabis, induces a toxicological effect on brain which renders the individual more sensitive to reinforcing effects of harder drugs. Evidence for this theory is supported by controlled animal model intervention studies (e.g., (Collins & Izenwasser, 2004)). Animal studies on the effects of caffeine intake on increasing later alcohol consumption (Hou et al., 2016; Kunin et al., 2000; SanMiguel et al., 2019) supports that the gateway hypothesis could also be applied to the association between caffeine and alcohol. Thus, one possible explanation of higher alcohol sipping rate of the daily-drinking group in our study is that the substances contained in caffeinated soda (caffeine and sugar) may have induced neurophysiological effects and reinforced regular soda drinkers to try alcohol after 12 months.

Conversely, the “common liability” hypothesis posits that the progression from softer to harder substances results primarily from the intersection of a genetically-regulated under-controlled temperament with progressively expanding *access* to harder substances with advancement into young adulthood (Vanyukov et al., 2012). Cross-sectional observations supported the common liability hypothesis, as children and adolescents with disruptive behavior disorders such as conduct disorder, which is strongly linked to SUDs, showed deficits in executive function (e.g., impaired impulse control) (Matthys et al., 2013). Longitudinal studies also support the common liability hypothesis (Debenham et al., 2021), wherein impaired impulse control and high sensation-seeking in young adolescents are predictive of increased drinking over time (Castellanos-Ryan et al., 2011). Similarly, caffeinated soda intake itself may not directly cause children to initiate substance use, but the drinking behavior of the beverages could indicate high impulsivity, which may be linked to a high risk of initiating substance use in the near future, consistent with the common liability hypothesis (Vanyukov et al., 2012). Because caffeinated soda contains two addictive substances, sugar and caffeine, it is somewhat natural for children to prefer and seek the beverages (Temple, 2009). While this taste preference could lead to a seeking behavior and habitual consumption, previous studies have shown impulsivity as one of the most common traits of SUDs (Cyders et al., 2009; Magid & Colder, 2007). Therefore, the behavior of frequently consuming caffeinated soda could indicate a high risk of initiating substance use in the future, due to the common risk factors (e.g., high impulsivity) between the two behaviors, although further research is needed to disentangle this complex relationship.

To our knowledge, this is the first study to investigate the direct link between caffeinated soda intake in earlier childhood and the risk of substance use. The results are consistent with studies on caffeine consumption during adolescence and its association with future substance misuse (Barrense-Dias et al., 2016; Leal & Jackson, 2019), supporting a higher risk of caffeinated soda consumption in childhood, particularly regarding vulnerability to future substance misuse. Such information is invaluable, as caffeinated soda is incomparably the most common medium for caffeine consumption in childhood, and the risk of substance misuse should be detected before adolescence, the most common period of substance use onset.

Our results have important implications for public health recommendations, as our study provides novel insight into the neurobehavioral correlates of caffeinated soda consumption in children, which has rarely been evaluated. At the same time, a few limitations of our research and future directions for further investigations should be discussed. First, we did not perform functional connectivity analyses or multivoxel pattern analyses, which may have provided additional insight into the effects of caffeinated soda intake. Second, as the 9–10-year-old children in this study had not yet started other substances except for alcohol, such as tobacco or marijuana, future work using the longitudinal 10-year follow-up data of the ABCD study should examine whether frequent consumption of caffeinated soda is associated with alcohol or other substance misuse. Third, we acknowledge that multiple variables other than caffeinated soda intake may mediate the relationship between neurobehavioral risk factors and future alcohol use; thus, extensive investigation of how caffeinated soda intake interacts with other SUD risk/protective factors is needed in the future. In addition, the effects of acute caffeine intake could have influenced the task performance of the soda drinking groups (Graczyk et al., 2018). Thus, future studies investigating the longitudinal effects of caffeinated soda intake should control for the acute caffeine consumption of children. Lastly, the ABCD dataset included only a small set of measures of food/drink consumption. We relied on a self-report measure consisting of a single item to assess caffeinated soda intake. For a more comprehensive understanding of the risks associated with caffeinated soda intake, it is critical for future research to integrate more objective and detailed assessments of soda consumption. Moreover, it is essential to differentiate the effects of caffeine and sweeteners by including more diverse measures of caffeine or sugar consumption.

In conclusion, our results revealed the potential risks of caffeinated soda consumption in children by investigating the associations between caffeinated soda consumption and risk factors for SUDs and examining the ability of caffeinated soda consumption to predict future alcohol sipping, using the large ABCD dataset. While previous research on the side effects of caffeinated soda consumption has been limited to negative physical consequences, the present results strongly suggest that caffeinated soda drinking in children is also associated with altered neurobehavioral function and can predict alcohol sipping after 12 months. Our study further suggests a strong need to develop evidence-based recommendations for caffeinated soda consumption in minors (Temple, 2018), as there is no consensus on a safe dose of caffeinated soda in children, and some children are at higher risk of adverse events from frequent caffeinated soda intake. Further clarification on the causal relationships and neuro-developmental evidence are needed to determine whether caffeinated soda is a warning sign for future substance misuse and whether it induces neurobehavioral impairments in children.

## Acknowledgements

We thank the families and children who participated, and continue to participate, in the ABCD study, as well as staff at the study sites who are involved in data collection and curating the data release. Data used in the preparation of this article were obtained from the Adolescent Brain Cognitive Development (ABCD) Study (https://abcdstudy.org), held in the NIMH Data Archive (NDA). This is a multisite, longitudinal study designed to recruit more than 10,000 children age 9-10 and follow them over 10 years into early adulthood. The ABCD Study® is supported by the National Institutes of Health and additional federal partners under award numbers U01DA041048, U01DA050989, U01DA051016, U01DA041022, U01DA051018, U01DA051037, U01DA050987, U01DA041174, U01DA041106, U01DA041117, U01DA041028, U01DA041134, U01DA050988, U01DA051039, U01DA041156, U01DA041025, U01DA041120, U01DA051038, U01DA041148, U01DA041093, U01DA041089, U24DA041123, U24DA041147. A full list of supporters is available at https://abcdstudy.org/federal-partners.html. A listing of participating sites and a complete listing of the study investigators can be found at https://abcdstudy.org/consortium_members/. ABCD consortium investigators designed and implemented the study and/or provided data but did not necessarily participate in analysis or writing of this report. This manuscript reflects the views of the authors and may not reflect the opinions or views of the NIH or ABCD consortium investigators. The ABCD data repository grows and changes over time. The ABCD data used in this report came from [NIMH Data Archive Digital Object Identifier (10.15154/1503209)]. DOIs can be found at http://dx.doi.org/10.15154/1503209.

