## Supplementary materials for "Caffeinated soda intake in children is associated with neurobehavioral risk factors for substance misuse"

### Working Memory

List Sorting Working Memory Test (list sorting test; Tulsky et al., 2013) requires participants to sort and sequence visual and auditory stimuli. For the Dimensional Change Card sort Test (card sort test; Tulsky et al., 2013), participants were asked to sort cards according to either color (when the target card has a contour) or shape (when the card without a contour is shown).

The emotional N-back (EN-back) task (Cohen et al., 2015) consists of two different blocks: the 0-back block requires participants to respond when the current stimulus is the same as a target, and the 2-back block requires participants to respond when the current stimulus is the same as the one shown in the two previous trials. Due to a difference in the working memory (WM) load between the two blocks, accuracy during the 2-back block is commonly used as a behavioral measure of WM, and the 2-back versus 0-back condition contrast is commonly used to examine the neural correlates of WM (Casey et al., 2018) in the functional magnetic resonance imaging (fMRI) analysis. See Casey et al. (2018) for more details about the task and the ROIs.

### Impulsivity

The stop signal task (SST; Logan et al., 1984) requires participants to withhold motor responses to a "Go" stimulus when a "Stop" signal is unpredictably presented after a short delay. The delay between the "Go" stimulus and the "Stop" signal is termed the stop signal delay (SSD). The SSD was initially set to 50 ms. Following successful or unsuccessful inhibition, the SSD was increased or decreased by 50 ms to make the task more difficult or easier, respectively. Each run contained 180 trials, and each trial began with the “Go” cue. Participants were instructed to respond to the cue as quickly and accurately as possible. Thirty trials from each run (16.67%) presented a "Stop" signal, which was an upward arrow displayed for 300 ms. Stop signal reaction time (SSRT) was used as a behavioral measure of impulsivity, calculated by subtracting the mean SSD from the mean “Go” response time (Logan et al., 1984); a shorter SSRT means better response inhibition. A “correct stop versus correct go” contrast was used for the fMRI analysis. More information about the SST is available in Casey et al. (2018).

**Reward processing**

Cash choice task asked the following single-item question: “Let’s pretend a kind person wanted to give you some money. Would you rather have $75 in 3 days or $115 in 3 months?” The former choice reflects poor delayed gratification, with a higher delay discounting rate, while the latter choice implies a smaller delay discounting rate. Responses were scored as 1 ($75 in 3 days) or 2 ($115 in 3 months).

Each trial of the monetary incentive delay (MID) task (Knutson et al., 2000) begins with an incentive cue of five possible trial types (win $20, win $5, lose $0.2, lose $5, and $0 [no money at stake]), followed by a delay (1,500–4,000 ms) and presentation of a target, to which the participant must respond while the target is rendered on the screen to either win money or avoid losing money. The trial ends with feedback on whether the participant won or lost money. Each participant performed 40 reward anticipation, 40 loss anticipation, and 20 no money anticipation trials, resulting in a total of 100 trials. The difference in hit rates between the reward and neutral trials indicates motivation to win the monetary reward.

**MRI protocol**

Imaging-derived phenotype scores provided by the Adolescent Brain Cognitive Development (ABCD) study were used. The data acquisition protocol was matched for the three 3T scanners used (Siemens Prisma, General Electric 750, and Philips). We analyzed beta weights within each ROI for the 2-back vs. 0-back contrast on the EN-back task, the correct stop vs. correct go contrast on the SST, and the reward vs. neutral anticipation contrast on the MID task. More details about the fMRI imaging protocols, such as the scanning parameters in the ABCD study, have been reported previously (Casey et al., 2018).

**Confounding variables**

Socioeconomic status (SES) was estimated using family income and parental education reported by the parents. Family history of substance use was measured using a parental yes/no response to the following question: "Has any blood relative of your child ever had any problems due to drugs, such as: marital separation or divorce; fired from work; arrests or driving under influence; adverse health effects; in a drug treatment program; Suspended or expelled from school two or more times; Isolated oneself from the family, caused arguments, or often intoxicated." The same question was asked to measure the family history of alcohol use, except that "drug" was replaced with "alcohol".

Parental monitoring was measured using the ABCD Parental Monitoring Survey, in which children answered five questions (“How often do your parents/guardians know where you are?”, “How often do your parents know who you are with, or when you are not at school and away from home?”, “If you are at home alone, do you know how to get in touch with your parents or guardians?”, “How often do you talk to your parent or guardian about your plans for the coming day, such as what will happen at school or what you are going to do with friends?”, “In an average week, how many times do you eat dinner with your parents/guardians?”). Responses ranged from 1 (never) to 5 (always or almost always), and the mean of all five responses was used for the analysis. SES is measured as the total combined family income and the parents’ highest education level. Parents’ education level was determined as the average of both parents’ levels, or that of the responding parent in the case of single parents.

Sleep deprivation was measured using the ABCD Parent Sleep Disturbance Scale for Children, for which parents responded to the question, “How many hours of sleep does your child get on most nights?” (1=9–11 hours, 2=8–9 hours, 3=7–8 hours, 4=5–7 hours, 5=less than 5 hours). Physical activity was measured using the ABCD Parent Sports and Activities Involvement Questionnaire, which asks whether the child has ever continuously participated in specific sports and activities, such as football, field hockey, climbing, or basketball, for ≥ 4 months. The number of “yes” responses to the 29 questions was summed for the analysis. Body mass index (BMI) was calculated according to the children’s body weight and height.

Externalizing and internalizing behaviors were measured using the parental report of the Child Behavior Checklist (Achenbach, 2009), which comprises 113 items about child behavior over the past 6 months. Internalizing and externalizing scores were measured based on the following syndrome scores: anxious/depressed, withdrawn/depressed, somatic complaints, social problems, thought problems, rule-breaking behavior, and aggressive behavior.

The fMRI data were collected from 22 different sites in the United States using three different MRI scanners (Siemens, General Electric, and Philips). All data were dummy coded and controlled in further analyses to control the site effects and scanner effects.

**LASSO regression**

A least absolute shrinkage and selection operator (LASSO) regression offers automatic variable selection via a penalized least squares method, which increases the interpretability of the findings, particularly when the multiple variables share a common characteristic to some extent. For example, each task measure (e.g., the EN-back task, list sorting test, and card sort test) within the cognitive risk factors is often moderately correlated with each other because they share common cognitive processing (e.g., WM). In this case, LASSO regression shrinks coefficients of variables with less explanatory power to zero and selects only variables with the highest power for further analyses (Tibshirani, 1996).

We used the R package called "easyml" (v0.1.1), which is a toolkit to easily build and evaluate machine learning models using the glmnet package (Friedman et al., 2010) for LASSO regression and several other machine learning algorithms (https://github.com/CCS-Lab/easyml; Ahn et al., 2017). The mixing parameter alpha was set to 1 to conduct the LASSO (1 for LASSO, 0 for ridge, and 0–1 for the elastic net). We randomly allocated half of the participants into training set and the other half into test set (i.e., train-test split = 0.5). To ensure the robustness of the results, accounting for potential variations in training and test set divisions, we performed random splitting a thousand times and evaluated model performance for each train-test split (see Ahn et al. (2017) for more details). Given that previous results have revealed that the elastic net often performs better for highly correlated predictors than for LASSO regression (Zou & Hastie, 2005), we tested the elastic net by setting alpha to a value between 0 and 1. However, there were no significant differences between the LASSO and elastic net results; thus, we only report the LASSO results.

**Additional analysis #1**

To thoroughly investigate the link between caffeinated soda consumption and future alcohol sipping, we used a continuous measure of caffeinated soda intake instead of the dichotomous daily- and non-drinkers classification. By using data from 4,517 participants (see **Fig. S1**), we performed the binomial LASSO regression analysis as described in the main manuscript (**Caffeinated soda intake and future alcohol sipping**), but with a continuous measure for the caffeinated soda intake. As shown in **Fig. S4**, the results support the main finding of an association between caffeinated soda intake and future alcohol sipping (**Fig. 3A**), as continuous soda intake was identified as one of the predictors that distinguished alcohol sipping after 12 months (mean estimate of coefficients=0.024, 95% CI=[0.003, 0.046]). The model had mean AUC values of 0.81 and 0.75 for the training and test sets, respectively (**Fig. S4B**).

**Additional analysis #2**

To include as many participants as possible, we opted to use behavioral measures of risk factors to predict future alcohol sipping. Thus, we performed the binomial LASSO regression analysis once again, using data from 8,939 participants without any missing data pertaining to soda intake, alcohol sipping, and all confounding variables. The results (**Fig. S5**) confirm the main result of the study, which is the link between caffeinated soda intake and future alcohol sipping (**Fig. 3A**). Even in the larger sample, the continuous soda intake was identified as one of the predictors that distinguished alcohol sipping after 12 months (mean estimate of coefficients=1.941, 95% CI=[1.935, 1.947]). The model had mean AUC values of 0.8 and 0.79 for the training and test sets, respectively (**Fig. S5B**).

**Additional analysis #3**

Using the largest sample of 8,939 participants, we employed hierarchical logistic regression analysis to verify whether baseline continuous measurements of caffeinated soda intake can anticipate alcohol sipping after 12 months. We used the "lm" function in R and tested whether caffeinated soda intake remains as a significant predictor even after controlling for all confounding variables, without including neural data (task-based fMRI data). The main predictor of interest was continuous measure of caffeinated soda intake. To ensure that the main predictor accounted for more variance than the confounding variables, we set a three-level hierarchical model (**Table S4**). The first level was the predictors as the confounding variables, including sex, age, racial identities, family income, parental education, physical activity, parental monitoring, family history of drug and alcohol abuse, sleep deprivation, externalizing and internalizing behaviors, and data collection sites. At the second level, we included alcohol sipping at baseline as an additional predictor. Finally, at the third level, continuous measure of caffeinated soda intake was added as a predictor to examine the effect of caffeinated soda intake. In other words, we first estimated variances explained by the variables in the first level, then we added baseline alcohol sipping in the second level to explain additional variance beyond that explained by the variables in the first level. Lastly, we examined whether caffeinated soda intake could significantly explain additional variance beyond that explained by the variables in the second level.

Two conditions must be satisfied to conclude that caffeinated soda intake is responsible for group differences in future alcohol sipping. First, we conducted analysis of variance (ANOVA) between the third-level analysis and the other two lower-level analyses; the third level should show higher significance than the previous levels. Additionally, the categorical caffeinated soda intake regressor should be significant ($p<0.05$) in the third-level analysis. We evaluated these conditions to determine whether caffeinated soda intake is a significant predictor of alcohol sipping after 12 months.

The adjusted R-square value of Model 3 revealed a significant difference compared with Model 1 ($F(2, 8{,}902)=356.50$, $p<2.20\text{e-}16$) and Model 2 ($F(1{,}8{,}902)=13.02$, $p<0.001$), supporting the main findings of the link between caffeinated soda intake and future alcohol sipping. Higher caffeinated soda consumption significantly explained alcohol sipping after 12 months ($\beta=0.004$, 95% CI=[0.002,

0.006]) even after controlling for confounding variables and baseline alcohol sipping (**Table S4**).

**Figure S1:** Flowchart showing the process of inclusion and exclusion of participants.

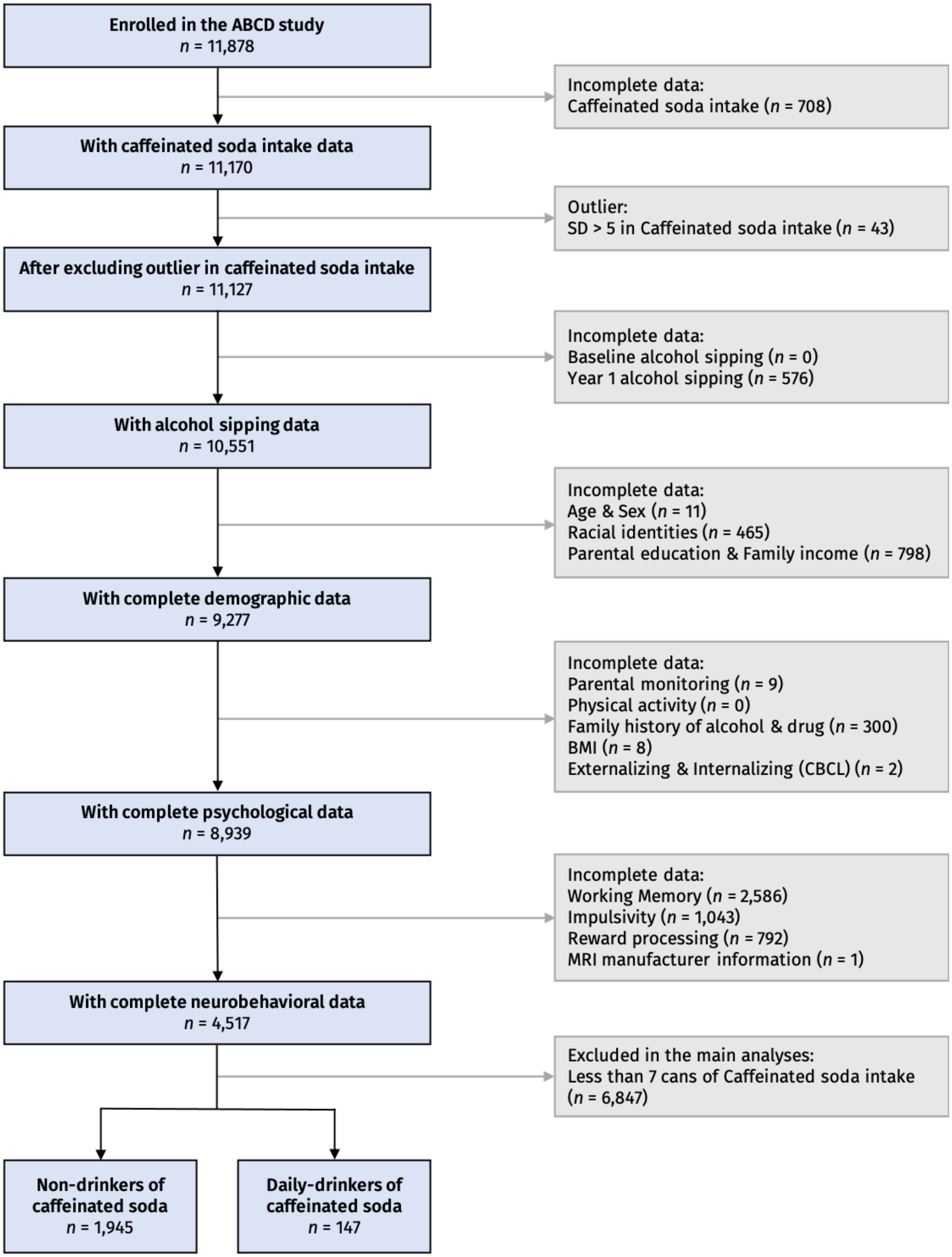

**Figure S2: Related to Figure 2.** Results of the LASSO regression predicting daily caffeinated soda intake (daily-drinking group vs. non-drinking group)

While **Figure 2** only shows survived variables without site variables, this figure contains all independent variables used for the prediction, including site variables and variables that did not survive. Mean of coefficient estimates for survived variables are shown in red (positive coefficients) or green (negative coefficients) dots. Each error bar indicates 95% confidence interval. Variables that did not survive (i.e., 95% confidence interval includes zero) are shown in grey dots.

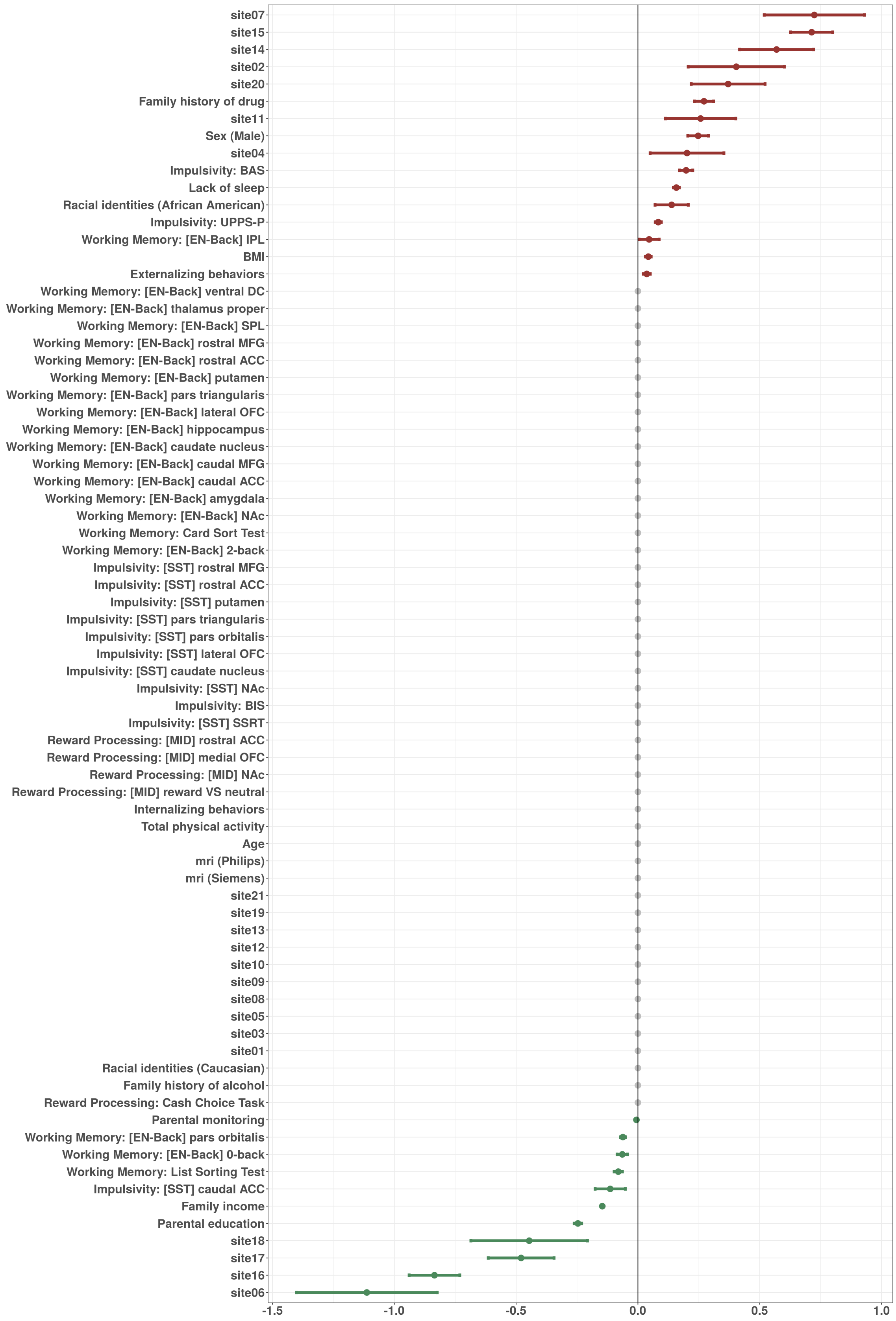
site07
site15
site14
site02
site20
Family history of drug
site11
Sex (Male)
site04
Impulsivity: BAS
Lack of sleep
Racial identities (African American)
Impulsivity: UPPS-P
Working Memory: [EN-Back] IPL
BMI
Externalizing behaviors
Working Memory: [EN-Back] ventral DC
Working Memory: [EN-Back] thalamus proper
Working Memory: [EN-Back] SPL
Working Memory: [EN-Back] rostral MFG
Working Memory: [EN-Back] rostral ACC
Working Memory: [EN-Back] putamen
Working Memory: [EN-Back] pars triangularis
Working Memory: [EN-Back] lateral OFC
Working Memory: [EN-Back] hippocampus
Working Memory: [EN-Back] caudate nucleus
Working Memory: [EN-Back] caudal MFG
Working Memory: [EN-Back] caudal ACC
Working Memory: [EN-Back] amygdala
Working Memory: [EN-Back] NAc
Working Memory: Card Sort Test
Working Memory: [EN-Back] 2-back
Impulsivity: [SST] rostral MFG
Impulsivity: [SST] rostral ACC
Impulsivity: [SST] putamen
Impulsivity: [SST] pars triangularis
Impulsivity: [SST] pars orbitalis
Impulsivity: [SST] lateral OFC
Impulsivity: [SST] caudate nucleus
Impulsivity: [SST] NAc
Impulsivity: BIS
Impulsivity: [SST] SSRT
Reward Processing: [MID] rostral ACC
Reward Processing: [MID] medial OFC
Reward Processing: [MID] NAc
Reward Processing: [MID] reward VS neutral
Internalizing behaviors
Total physical activity
Age
mri (Philips)
mri (Siemens)
site21
site19
site13
site12
site10
site09
site08
site05
site03
site01
Racial identities (Caucasian)
Family history of alcohol
Reward Processing: Cash Choice Task
Parental monitoring
Working Memory: [EN-Back] pars orbitalis
Working Memory: [EN-Back] 0-back
Working Memory: List Sorting Test
Impulsivity: [SST] caudal ACC
Family income
Parental education
site18
site17
site16
site06
-1.5
-1.0
-0.5
0.0
0.5
1.0

**Figure S3: Related to Figure 3.** Results of the LASSO regression predicting alcohol sipping after 12 months

While **Figure 3** only shows survived variables without site variables, this figure contains all independent variables used for the prediction, including site variables and variables that did not survive. Mean of coefficient estimates for survived variables are shown in red (positive coefficients) or green (negative coefficients) dots. The error bar indicates 95% confidence interval. Variables that did not survive (i.e., 95% confidence interval includes zero) are shown in grey dots.

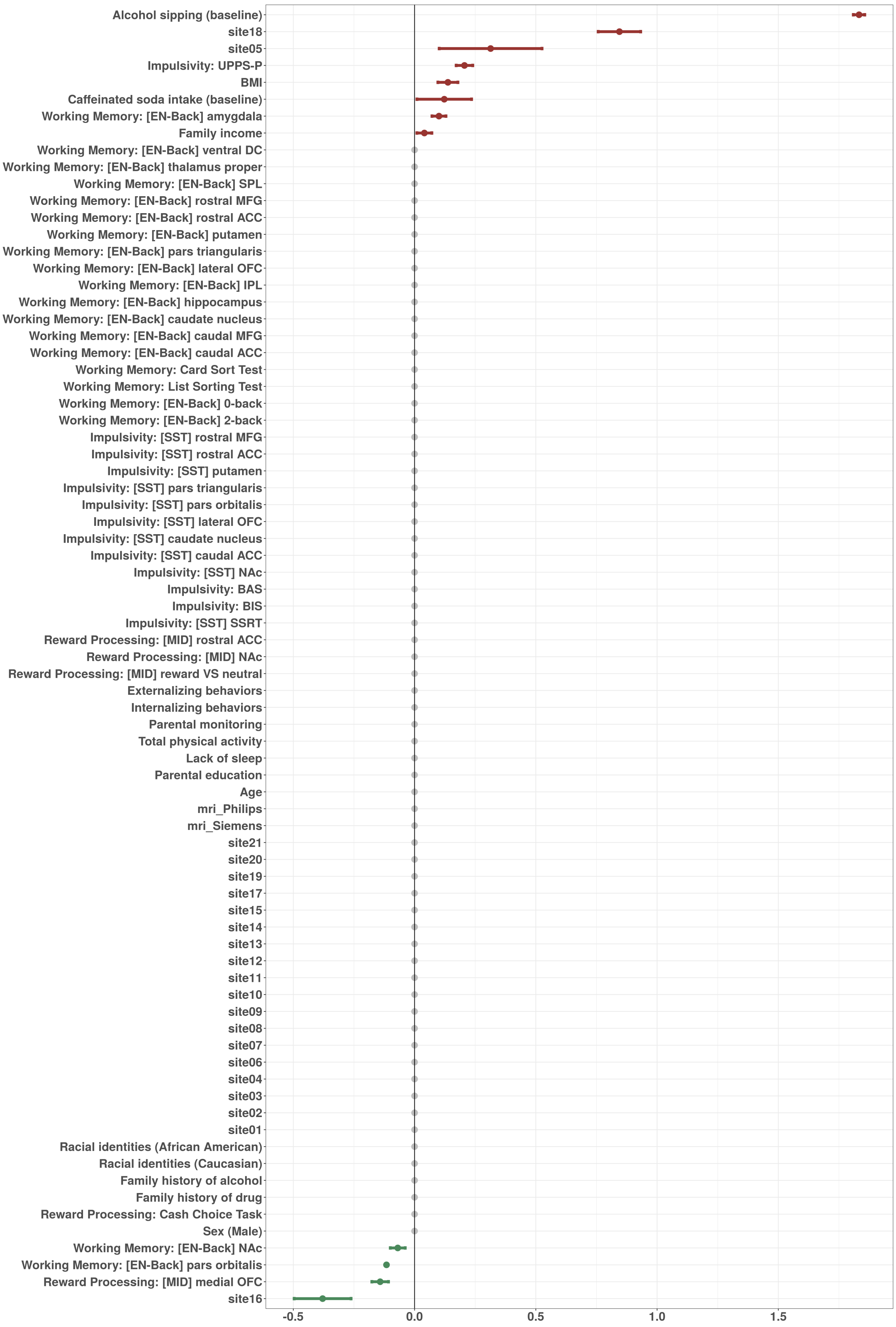
Alcohol sipping (baseline)
site18
site05
Impulsivity: UPPS-P
BMI
Caffeinated soda intake (baseline)
Working Memory: [EN-Back] amygdala
Family income
Working Memory: [EN-Back] ventral DC
Working Memory: [EN-Back] thalamus proper
Working Memory: [EN-Back] SPL
Working Memory: [EN-Back] rostral MFG
Working Memory: [EN-Back] rostral ACC
Working Memory: [EN-Back] putamen
Working Memory: [EN-Back] pars triangularis
Working Memory: [EN-Back] lateral OFC
Working Memory: [EN-Back] IPL
Working Memory: [EN-Back] hippocampus
Working Memory: [EN-Back] caudate nucleus
Working Memory: [EN-Back] caudal MFG
Working Memory: [EN-Back] caudal ACC
Working Memory: Card Sort Test
Working Memory: List Sorting Test
Working Memory: [EN-Back] 0-back
Working Memory: [EN-Back] 2-back
Impulsivity: [SST] rostral MFG
Impulsivity: [SST] rostral ACC
Impulsivity: [SST] putamen
Impulsivity: [SST] pars triangularis
Impulsivity: [SST] pars orbitalis
Impulsivity: [SST] lateral OFC
Impulsivity: [SST] caudate nucleus
Impulsivity: [SST] caudal ACC
Impulsivity: [SST] NAc
Impulsivity: BAS
Impulsivity: BIS
Impulsivity: [SST] SSRT
Reward Processing: [MID] rostral ACC
Reward Processing: [MID] NAc
Reward Processing: [MID] reward VS neutral
Externalizing behaviors
Internalizing behaviors
Parental monitoring
Total physical activity
Lack of sleep
Parental education
Age
mri_Philips
mri_Siemens
site21
site20
site19
site17
site15
site14
site13
site12
site11
site10
site09
site08
site07
site06
site04
site03
site02
site01
Racial identities (African American)
Racial identities (Caucasian)
Family history of alcohol
Family history of drug
Reward Processing: Cash Choice Task
Sex (Male)
Working Memory: [EN-Back] NAc
Working Memory: [EN-Back] pars orbitalis
Reward Processing: [MID] medial OFC
site16
-0.5
0.0
0.5
1.0
1.5

1

**Figure S4:** Results of LASSO regression predicting alcohol sipping after 12 months (using continuous measure of caffeinated soda intake; $N = 4{,}529$)

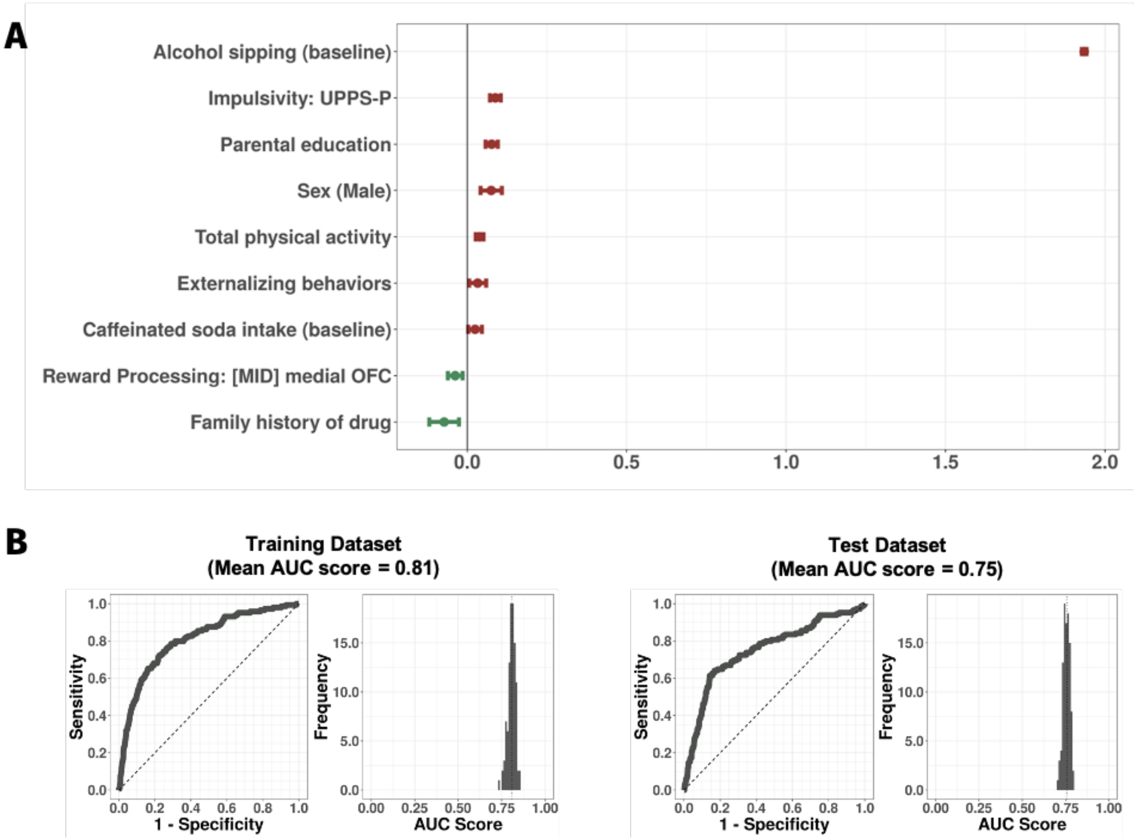


**A.** Estimates of coefficients of the survived variables (x axis: coefficient estimates; y axis: predictor). The site variables were excluded for clarification. Each dot indicates mean of the coefficient, positive coefficient in red and negative coefficient in green. Each error bar indicates 95% confidence interval. **B.** Distribution of the area under the curve (AUC) values (left) and a representative receiver-operation characteristic (ROC) curve (right) for the training and test datasets. *Abbreviations*. LASSO, least absolute shrinkage and selection operator; UPPS-P, Urgency-Premeditation-Perseverance-Sensation Seeking-Positive Urgency; MID, Monetary Incentive Delay Task; OFC, Orbitofrontal cortex.

**Figure S5:** Results of LASSO regression predicting alcohol sipping after 12 months (using continuous measure of caffeinated soda intake and using only behavioral measures without fMRI measures; $N = 8{,}939$)

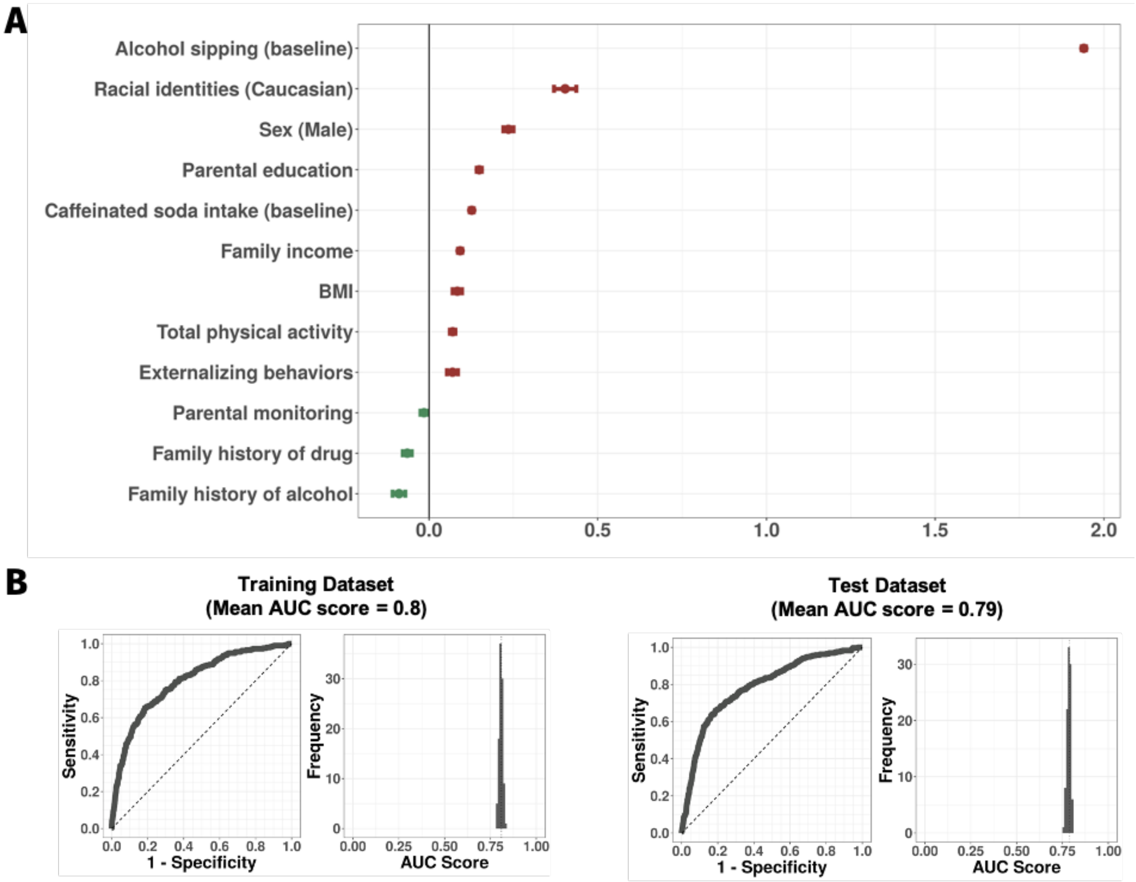


**A.** Estimates of coefficients of the survived variables (x axis: coefficient estimates; y axis: predictor). The site variables were excluded for clarification. Each dot indicates mean of the coefficient, positive coefficient in red and negative coefficient in green. Each error bar indicates 95% confidence interval. **B.** Distribution of the area under the curve (AUC) values (left) and a representative receiver-operation characteristic (ROC) curve (right) for the training and test datasets. *Abbreviations*. LASSO, least absolute shrinkage and selection operator; BMI, body mass index.

**Table S1.** Characteristics of included and excluded samples

| | Included | Excluded | P-value | Effect size |
|---|---|---|---|---|
| **Sample size** | 2092 (23.4%) | 6847 (76.6%) | | |
| **Soda intake/week** | 0.78 (0.71, 0.85) | 1.05 (1.02, 1.07) | <0.001 | 0.10 |
| **Baseline alcohol sipping (%)** | 324 (15.49%) | 1305 (19.06%) | <0.001 | 0.04 |
| **Year 1 alcohol sipping (%)** | 90 (4.30%) | 492 (7.19%) | <0.001 | 0.05 |
| **Sex - Male (%)** | 954 (45.60%) | 3734 (54.53%) | <0.001 | 0.08 |
| **Racial identities - Caucasian (%)** | 1791 (85.61%) | 5225 (76.31%) | <0.001 | 0.10 |
| **Racial identities - African American (%)** | 154 (7.36%) | 1088 (15.89%) | <0.001 | 0.10 |
| **Age** | 9.49 (9.48, 9.50) | 9.49 (9.48, 9.49) | 0.965 | 0 |
| **Family income** | 8.01 (7.97, 8.05) | 7.32 (7.29, 7.35) | <0.001 | -0.32 |
| **Parental education** | 17.39 (17.35, 17.44) | 16.57 (16.54, 16.60) | <0.001 | -0.36 |
| **Parental monitoring** | 4.46 (4.45, 4.47) | 4.38 (4.38, 4.39) | <0.001 | -0.16 |
| **Family history - drug (%)** | 584 (27.92%) | 2205 (32.20%) | <0.001 | 0.04 |
| **Family history - alcohol (%)** | 915 (43.74%) | 3290 (48.05%) | <0.001 | 0.04 |
| **BMI** | 18.01 (17.93, 18.09) | 18.8 (18.75, 18.85) | <0.001 | 0.20 |
| **Physical activity** | 4.25 (4.20, 4.31) | 3.71 (3.68, 3.74) | <0.001 | -0.21 |
| **Lack of sleep** | 1.50 (1.48, 1.51) | 1.70 (1.69, 1.71) | <0.001 | 0.27 |
| **Externalizing behaviors** | 44.11 (43.91, 44.32) | 45.74 (45.62, 45.87) | <0.001 | 0.17 |
| **Internalizing behaviors** | 48.24 (48.02, 48.46) | 48.41 (48.28, 48.54) | 0.52 | 0.02 |

*Note.* Data are the means (95% confidence interval) or numbers (percentage) for each sample. Difference between the included and excluded samples was examined using a two-sample t-test or a chi-square test. Effect size for the two-sample t-test was estimated using Cohen's $d$ ($d$ = 0.2, small; $d$ = 0.5, medium; $d$ = 0.8, large effect size). Effect size for the chi-square test was estimated using Cramer's V (when degrees of freedom = 1, V = 0.1, small; V = 0.3, medium; V = 0.5, large). *Abbreviations.* BMI, Body Mass Index.

**Table S2.** Characteristics of the participants

| | Daily-drinking (≥ 7 can/week) | Non-drinking (0 can) | P-value | Effect size |
|---|---|---|---|---|
| **Sample size** | 147 (7.03%) | 1945 (92.97%) | | |
| **Soda intake/week** | 11.11 (10.71, 11.51) | 0 (0, 0) | <0.001 | 3.25 |
| **Baseline alcohol sipping (%)** | 32 (21.77%) | 292 (15.01%) | 0.029 | 0.05 |
| **Year 1 alcohol sipping (%)** | 12 (8.16%) | 78 (4.01%) | 0.017 | 0.05 |
| **Sex - Male (%)** | 82 (55.78%) | 872 (44.83%) | 0.010 | 0.06 |
| **Racial identities - Caucasian (%)** | 105 (71.43%) | 1686 (86.68%) | <0.001 | 0.11 |
| **Racial identities - African American (%)** | 30 (20.41%) | 124 (6.38%) | <0.001 | 0.14 |
| **Age** | 9.54 (9.50, 9.58) | 9.48 (9.47, 9.49) | 0.211 | 0.11 |
| **Family income** | 6.86 (6.65, 7.06) | 8.10 (8.06, 8.14) | <0.001 | -0.57 |
| **Parental education** | 16.09 (15.88, 16.29) | 17.49 (17.44, 17.54) | <0.001 | -0.62 |
| **Parental monitoring** | 4.34 (4.29, 4.39) | 4.47 (4.46, 4.48) | <0.001 | -0.24 |
| **Family history - drug (%)** | 58 (39.46%) | 526 (27.04%) | 0.001 | 0.07 |
| **Family history - alcohol (%)** | 74 (50.34%) | 841 (43.24%) | 0.094 | 0.04 |
| **BMI** | 19.35 (18.94, 19.77) | 17.91 (17.83, 17.99) | <0.001 | 0.33 |
| **Physical activity** | 3.51 (3.31, 3.71) | 4.31 (4.25, 4.37) | <0.001 | -0.31 |
| **Lack of sleep** | 1.84 (1.76, 1.91) | 1.47 (1.46, 1.49) | <0.001 | 0.46 |
| **Externalizing behaviors** | 46.23 (45.3, 47.16) | 43.95 (43.74, 44.17) | 0.005 | 0.22 |
| **Internalizing behaviors** | 48.57 (47.69, 49.46) | 48.21 (47.99, 48.44) | 0.681 | 0.03 |
| **Working memory** | | | | |
| **List Sorting Test** | 99.65 (98.38, 100.92) | 104.51 (104.20, 104.83) | <0.001 | -0.33 |
| **Card sort test** | 96.18 (95.02, 97.35) | 99.89 (99.54, 100.24) | 0.005 | -0.25 |
| **[EN-Back] 2-back** | 0.79 (0.78, 0.80) | 0.81 (0.80, 0.81) | 0.045 | -0.17 |
| **[EN-Back] 0-back** | 0.85 (0.85, 0.86) | 0.88 (0.87, 0.88) | 0.002 | -0.25 |
| **Impulsivity** | | | | |
| **UPPS-P** | 42.06 (41.46, 42.66) | 39.81 (39.64, 39.97) | <0.001 | 0.31 |
| **BIS** | 9.46 (9.15, 9.77) | 9.33 (9.25, 9.41) | 0.665 | 0.04 |
| **BAS** | 22.26 (21.71, 22.81) | 19.46 (19.32, 19.61) | <0.001 | 0.42 |
| **[SST] SSRT** | 304.93 (299.57, 310.29) | 298.08 (296.58, 299.58) | 0.226 | 0.10 |
| **Reward processing** | | | | |
| **Cash Choice Task (%)** | 88 (59.86%) | 1184 (60.87%) | 0.809 | 0.01 |
| **[MID] reward vs. neutral** | 0.11 (0.10, 0.12) | 0.12 (0.11, 0.12) | 0.461 | -0.06 |

*Note.* Data are the means (95% confidence interval) for each group (daily-drinking and non-drinking group) or numbers (percentage) for sample size, sex, racial identities, family history, and cash choice task. Difference between the daily-drinking group and non-drinking was examined using a two-sample t-test or a chi-square test. Effect size for the two-sample t-test was estimated using Cohen's $d$ ($d$ = 0.2, small; $d$ = 0.5, medium; $d$ = 0.8, large effect size). Effect size for the chi-square test was estimated using Cramer's V (when degrees of freedom = 1, V = 0.1, small; V = 0.3, medium; V = 0.5, large). See **Table S3** for the beta coefficient estimates of the fMRI data. *Abbreviations.* BMI, Body Mass Index; BIS, Behavioral Inhibition System; BAS, Behavioral Activation System;

EN-back, Emotional N-Back Task; SST, Stop Signal Task; MID, Monetary Incentive Delay Task; SSRT, Stop Signal Reaction Time.

**Table S3.** Beta coefficient estimates for the fMRI data

| | Daily-drinking (≥ 7 can/week) | Non-drinking (0 can) | P-value | Effect size |
|---|---|---|---|---|
| **Sample size** | 147 (7.03%) | 1945 (92.97%) | | |
| **Working memory** | | | | |
| **[EN-Back] NAc** | -0.03 (-0.05, 0.00) | -0.04 (-0.05, -0.03) | 0.673 | -0.04 |
| **[EN-Back] amygdala** | -0.07 (-0.10, -0.04) | -0.07 (-0.07, -0.06) | 0.875 | 0.01 |
| **[EN-Back] caudal ACC** | 0.04 (0.02, 0.06) | 0.05 (0.04, 0.05) | 0.652 | 0.04 |
| **[EN-Back] caudal MFG** | 0.12 (0.10, 0.15) | 0.13 (0.12, 0.13) | 0.869 | 0.01 |
| **[EN-Back] caudate nucleus** | 0.05 (0.03, 0.08) | 0.04 (0.03, 0.04) | 0.387 | -0.07 |
| **[EN-Back] hippocampus** | -0.06 (-0.07, -0.04) | -0.07 (-0.08, -0.07) | 0.352 | -0.07 |
| **[EN-Back] IPL** | 0.06 (0.04, 0.08) | 0.05 (0.05, 0.06) | 0.569 | -0.05 |
| **[EN-Back] lateral OFC** | -0.08 (-0.16, -0.01) | -0.02 (-0.04, -0.01) | 0.241 | 0.08 |
| **[EN-Back] pars orbitalis** | -0.20 (-0.29, -0.10) | -0.07 (-0.09, -0.05) | 0.071 | 0.13 |
| **[EN-Back] pars triangularis** | -0.08 (-0.12, -0.03) | -0.06 (-0.07, -0.05) | 0.591 | 0.04 |
| **[EN-Back] putamen** | -0.01 (-0.03, 0.01) | -0.02 (-0.03, -0.02) | 0.523 | -0.05 |
| **[EN-Back] rostral ACC** | -0.10 (-0.14, -0.07) | -0.11 (-0.11, -0.10) | 0.962 | 0.00 |
| **[EN-Back] rostral MFG** | 0.02 (-0.03, 0.07) | 0.09 (0.08, 0.09) | 0.043 | 0.14 |
| **[EN-Back] SPL** | 0.04 (0.02, 0.06) | 0.04 (0.03, 0.04) | 0.839 | -0.02 |
| **[EN-Back] thalamus proper** | 0.05 (0.03, 0.06) | 0.04 (0.03, 0.04) | 0.491 | -0.06 |
| **[EN-Back] ventral DC** | 0.05 (0.02, 0.07) | 0.04 (0.04, 0.05) | 0.812 | -0.02 |
| **Impulsivity** | | | | |
| **[SST] NAc** | 0.01 (0.00, 0.03) | 0.01 (0.01, 0.01) | 0.814 | -0.02 |
| **[SST] caudal ACC** | 0.04 (0.03, 0.05) | 0.07 (0.06, 0.07) | 0.047 | 0.17 |
| **[SST] caudate nucleus** | 0.00 (-0.01, 0.02) | 0.03 (0.02, 0.03) | 0.098 | 0.14 |
| **[SST] lateral OFC** | 0.04 (0.01, 0.08) | 0.04 (0.03, 0.05) | 0.946 | -0.01 |
| **[SST] pars orbitalis** | 0.03 (-0.02, 0.07) | 0.00 (-0.01, 0.01) | 0.608 | -0.05 |
| **[SST] pars triangularis** | 0.05 (0.04, 0.07) | 0.05 (0.05, 0.05) | 0.823 | -0.02 |
| **[SST] putamen** | 0.00 (-0.01, 0.00) | 0.01 (0.01, 0.02) | 0.114 | 0.14 |
| **[SST] rostral ACC** | -0.02 (-0.04, 0.00) | -0.01 (-0.01, 0.00) | 0.464 | 0.06 |
| **[SST] rostral MFG** | 0.06 (0.05, 0.08) | 0.06 (0.06, 0.07) | 0.842 | -0.02 |
| **Reward processing** | | | | |
| **[MID] NAc** | 0.08 (0.06, 0.11) | 0.07 (0.07, 0.08) | 0.562 | -0.04 |
| **[MID] medial OFC** | 0.02 (-0.01, 0.06) | -0.02 (-0.03, -0.01) | 0.339 | -0.09 |
| **[MID] rostral ACC** | -0.01 (-0.03, 0.01) | 0.01 (0.01, 0.02) | 0.263 | 0.10 |

*Note* Data are the means (95% confidence interval) for each group (daily-drinking group and non-drinking group). Difference between the daily-drinking group and non-drinking was examined using a two-sample t-test. Effect size was estimated using Cohen's *d* (*d* = 0.2, small; *d* = 0.5, medium; *d* = 0.8, large effect size). *Abbreviations.* EN-back, Emotional N-Back Task; SST, Stop Signal Task; MID, Monetary Incentive Delay Task; NAc, Nucleus accumbens; ACC, Anterior Cingulate Cortex; MFG, Medial Frontal Gyrus; IPL, Inferior Parietal Lobule; OFC, Orbitofrontal Cortex; SPL, Superior Parietal Lobule; ventral DC, ventral diencephalon.

**Table S4.** Results of hierarchical logistic regression analysis predicting alcohol sipping after 12 months (using continuous measure of caffeinated soda intake and using only behavioral and demographic variables without fMRI measures; *N* = 8,939)

| | Model 1 | Model 2 | Model 3 |
|---|---|---|---|
| Sex (Male) | -0.021 (-0.031 - -0.010)*** | -0.016 (-0.026 - -0.006)** | -0.015 (-0.024 - -0.005)** |
| Racial identities (Caucasian) | 0.033 (0.012 - 0.053)** | 0.023 (0.003 - 0.043)* | 0.022 (0.002 - 0.042)* |
| Racial identities (African American) | 0.001 (-0.024 - 0.025) | 0.002 (-0.022 - 0.026) | 0.002 (-0.022 - 0.026) |
| Family income | 0.007 (0.000 - 0.014) | 0.004 (-0.003 - 0.011) | 0.005 (-0.002 - 0.012) |
| Parental education | 0.008 (0.002 - 0.015)* | 0.007 (0.000 - 0.014)* | 0.008 (0.001 - 0.014)* |
| Parental monitoring | -0.003 (-0.009 - 0.002) | -0.002 (-0.007 - 0.003) | -0.002 (-0.007 - 0.003) |
| Family history (drug) | -0.004 (-0.016 - 0.008) | -0.005 (-0.017 - 0.006) | -0.006 (-0.018 - 0.006) |
| Family history (alcohol) | -0.008 (-0.019 - 0.004) | -0.010 (-0.02 - 0.001) | -0.009 (-0.020 - 0.001) |
| BMI | 0.007 (0.002 - 0.012)* | 0.006 (0.001 - 0.011)* | 0.006 (0.001 - 0.011)* |
| Physical activity | 0.007 (0.001 - 0.012)* | 0.005 (0.000 - 0.011)* | 0.005 (0.000 - 0.011)* |
| Lack of sleep | 0.000 (-0.005 - 0.006) | 0.002 (-0.004 - 0.007) | 0.001 (-0.004 - 0.007) |
| Externalizing behaviors | 0.009 (0.003 - 0.015)** | 0.007 (0.001 - 0.013)* | 0.007 (0.001 - 0.013)* |
| Internalizing behaviors | -0.003 (-0.009 - 0.004) | -0.003 (-0.009 - 0.003) | -0.003 (-0.009 - 0.003) |
| Alcohol sipping (Baseline) | | 0.175 (0.162 - 0.188)*** | 0.175 (0.162 - 0.188)*** |
| Caffeinated soda intake (Baseline) | | | 0.004 (0.002 - 0.006)*** |

*Note.* The values are standardized logistic regression coefficients (95% confidence interval) from each step in the hierarchical analyses predicting alcohol sipping after 12 months. Significant variables predicting alcohol sipping are flagged with asterisks (*: $p<0.05$, **: $p<0.01$; ***: $p<0.001$). Higher consumption of caffeinated soda intake (Model 3) predicted a higher tendency for alcohol sipping after 12 months even after controlling for the confounding variables (13 variables from the top and 21 data collection sites in Model 1) and alcohol sipping collected at baseline (Model 2).
*Abbreviations.* BMI, Body Mass Index.
